# Alpha EEG power reflects the suppression of Pavlovian bias during social reinforcement learning

**DOI:** 10.1101/153668

**Authors:** James C Thompson, Margaret L Westwater

## Abstract

Socially appropriate behavior involves learning actions that are valued by others and those that have a social cost. Facial expressions are one way that others can signal the social value of our actions. The rewarding or aversive properties of signals such as smiles or frowns also evoke automatic approach or avoidance behaviors in receivers, and a Pavlovian system learns cues that predict rewarding or aversive outcomes. In this study, we examined the computational and neural mechanisms underlying interactions between Pavlovian and Instrumental systems during social reinforcement learning. We found that Pavlovian biases to approach cues predicting social reward and avoid cues predicting social punishment interfered with Instrumental learning from social feedback. While the computations underlying Pavlovian and Instrumental interactions remained the same as when learning from monetary feedback, Pavlovian biases from social outcomes to approach or withdraw were not significantly correlated with biases from money. Trial-by-trial measures of alpha (8-14Hz) EEG power was associated with suppression of Pavlovian bias to social outcomes, while suppression of bias from money was associated with theta (4-7Hz) EEG power. Our findings demonstrate how emotional reactions to feedback from others are balanced with the instrumental value of that feedback to guide social behavior.

**Significance statement:** A smile from another can be a signal to continue what we are doing, while an angry scowl is a sure sign to stop. Feedback from others such as this plays an important role in shapeing social behavior. The rewarding nature of a smile (or the aversive nature of a scowl) can also lead to automatic tendencies to approach (or avoid), and we can learn situations that predict positive or negative social outcomes. In this study, we examined the brain mechanisms that come into play when the instrumental demands of a situation are in conflict with our automatic biases to approach or withdraw, such as when we have to approach someone who is scowling at us or withdraw from someone who is smiling.

---

Learning socially appropriate behavior requires that we use feedback signals from others that indicate the value of our actions. Action value can be signaled by other people using facial expressions, such as a smile or a frown, and we can track social value information from past actions to help decide how we should act in the present (Frijda, 1986). Reinforcement learning (RL) approaches have proven useful in describing the action-outcome (Instrumental) learning process, including aspects of social learning in humans (Apps et al, 2016; Behrens et al., 2009; Lin et al., 2012; Zaki et al., 2016). At the same time that nonverbal social signals can guide action learning, the rewarding or aversive properties of these signals can generate affective responses and automatic behaviors, such as to approach (to a smile) or withdrawal (from a frown) (Fridja, 1986; Marsh et al., 2005). Responses to such events are controlled by a Pavlovian motivation system, which facilitates the learning of environmental cues predictive of reward or punishment, regardless of actions taken in response to such cues (Mackintosh, 1983). While instrumental and Pavlovian motivational systems might work towards the same goal of maximizing outcomes, these two systems can come into conflict; for example, we may need to approach someone who looks to be angry with us. Pavlovian-Instrumental (P-I) interactions have been demonstrated using nonsocial outcomes in humans (Talmi et al., 2008; Bray et al., 2008; Huys et al., 2011; Prevost et al., 2012; Geurts et al., 2013; Swart et al., 2017) and nonhuman animals (Rescorla & Solomon, 1967; Berridge, 1996). Balancing interactions between Instrumental and Pavlovian motivation systems during social behavior is an important component of self-control (Dayan et al., 2006). In this study we examined the neural and computational bases of the control of P-I interactions during learning from social feedback, and compared these to the neural and computational bases of the control of P-I interactions during learning from nonsocial feedback.

Medial and lateral prefrontal cortices (PFC) play a key role in controlling the balance between conflicting social, emotional, and cognitive demands (Etkin et al., 2006; Zaki et al., 2010). Oscillatory electro‑ and magnetoencephalographic (EEG & MEG) activity in the low-to-mid (1-30Hz) frequency range has been suggested to reflect top-down control (Helfrich & Knight, 2016; Buschman & Miller, 2014; von Stein & Sarnthein, 2000). Theta (4-7Hz) oscillations in medial and lateral PFC has been linked to adaptive responses to cognitive conflict and the need to engage control (Cavanagh & Frank, 2014). Alpha (8-14Hz) and beta (15-30Hz) oscillations have also been linked to inhibitory control and overriding automatic response tendencies (Hwang et al., 2014; Jensen and Mazaheri, 2010; Sadaghani & Kleinschmidt, 2016). Recently, it was shown that frontal theta (4-8Hz) EEG power reflected the trial-by-trial modulation of Pavlovian biases during instrumental learning based on monetary rewards and punishments (Cavanagh et al., 2013). Using a Go/No-Go task that orthogonalized action and outcome valence, combined with a computational model that incorporated Pavlovian and Instrumental learning, Cavanagh and colleagues (2013) found that greater midfrontal theta oscillations were associated with the successful suppression of Pavlovian biases during learning. Less is known about the neural mechanisms that underlie social-emotion control.

Here we examined if learning from social feedback used similar computational and neural mechanisms to learning from monetary feedback during P-I conflict. We developed a variant of the Go/No-Go learning task used by Cavanagh and colleagues (2013; Guitart-Masip et al., 2012) that used happy and angry facial expressions instead of monetary outcomes as feedback. We compared learning on a social Go/No-Go task to learning on the monetary Go/No-Go version of the task. EEG was recorded during task performance in order to examine neural activity associated with conflict between P-I systems. We then used a computational model that incorporated Pavlovian and Instrumental learning, based on that developed by Cavanagh and colleagues (2013), to determine on a trial-by-trial basis the neural mechanisms underlying the suppression of Pavlovian influence to social and monetary feedback.

## Materials & Methods

### Participants

Twenty-two healthy, right-handed adults (15 female; *M_age_* = 18.6 y; *SD =* 1.3; 13 White/Latino, 5 Asian, 2 African-American, 2 Biracial/Other) with normal or corrected-to-normal vision participated in the study. Participants were recruited as part of a larger study examining social functioning in late adolescence and early adulthood. Two participants were excluded as they were currently received psychological treatment, leaving a final N of 20. Prior to participation, participants provided written informed consent. The study procedure and all relevant materials were approved by the George Mason University Human Subjects Review Board (HSRB). Participants were informed at each session that they were free to withdraw from the study at any time for any reason, and they received monetary compensation ($15/hr plus winnings from monetary task) for their time.

### Experimental Design and Statistical Analysis

To examine interactions between Pavlovian and instrumental systems during social reinforcement learning, we adapted a Go/No-Go learning task (Figure 1) developed by Guitart-Masip and colleagues (2012). On each trial, participants were presented with one of four cue shapes for 1000ms, followed by variable interval (250-2000ms), and then a white circle (1000ms max) which acted as a response cue. After a 1000ms interval, participants then received outcomes for 2000ms. On win trials, 80% of correct actions to the response cues were rewarded and 20% received a neutral outcome, while 20% of incorrect actions were rewarded and 80% received a neutral outcome. For loss trials, 80% of correct actions received a neutral outcome and 20% received a punishment, while 80% of incorrect actions received a punishment and 20% received a neutral outcome. After a variable interval (1000-2000ms), a new trial was presented. On one half of trials, the correct action was to respond as quickly as possible (Go trials), and on the other half the correct action was to withhold response (No-Go trials). This design lead to four conditions, in two of which Pavlovian and Instrumental demands were not in conflict (Go to Win, No-Go to Avoid Loss), and two of which there was conflict between Pavlovian and Instrumental (Go to Avoid Less, No-Go to Win). There were 45 trials of each type for each version (monetary and social) of the task, with trial order randomized. Cue shapes were drawn from a set of eight shapes, split into a set of four for each task version, and shuffled across conditions for each participant.

**Figure 1.**
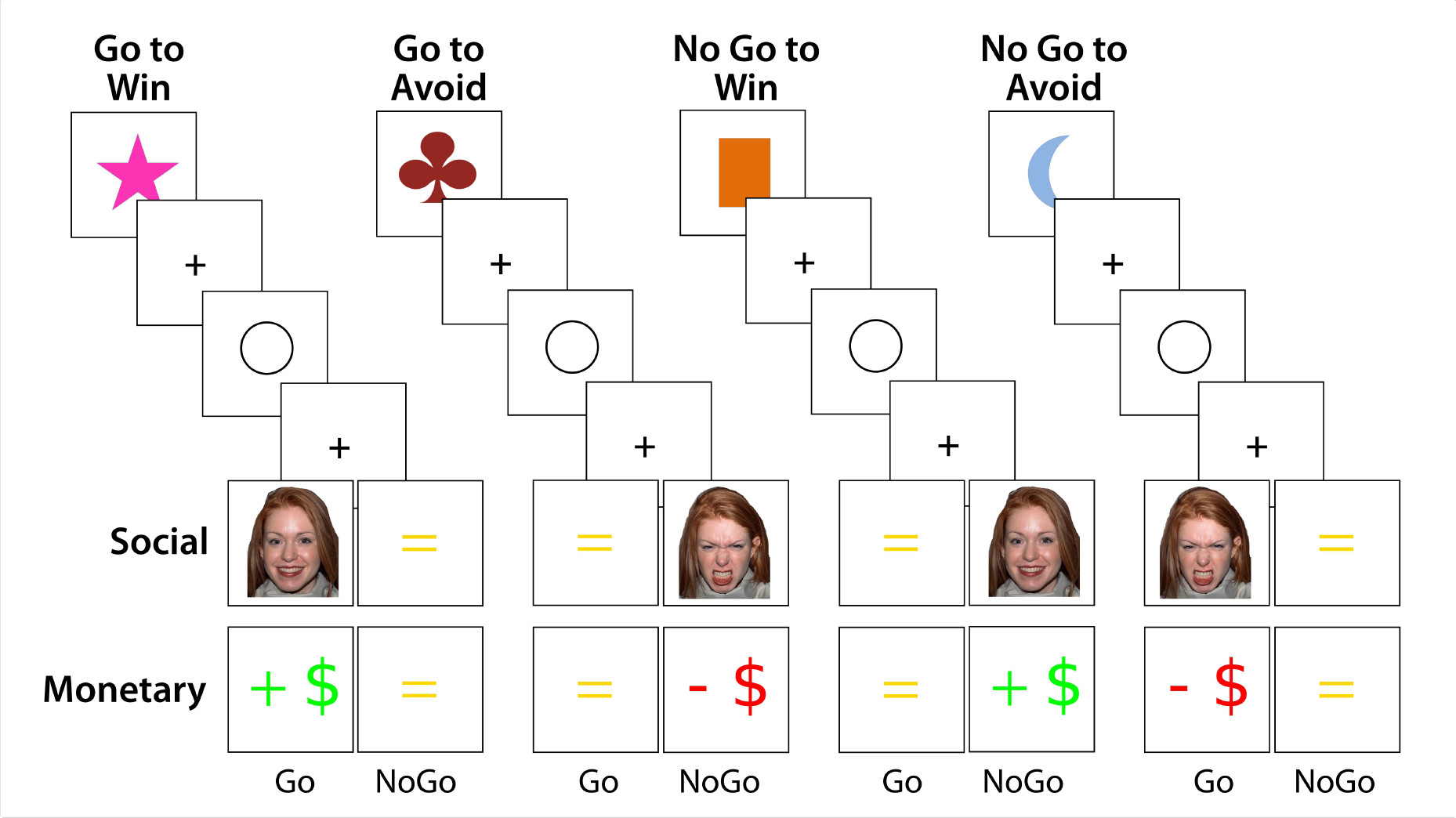
Task Design. Stimuli and trial design for the Social and Monetary Go/NoGo tasks. Task order was counterbalanced across participants.

In the monetary task, reward was indicated by a green “+ $”, indicating an increase in funds of 25c, punishment was indicated by a red “− $”, indicating a decrease in funds of 25c, or a yellow “=”, indicated a neutral outcome. In the social task, faces from the NimStim Set of Facial Expressions (Tottenham et al., 2009) were used as outcomes. Reward was indicated by a smiling (happy, mouth open) face, punishment was indicated by a frowning (angry) face, and neutral outcome was indicated by the same yellow “=” used in the monetary task. Eighteen male faces and eighteen female faces were used, with face identity/sex chosen at random on each trial. Order of task version presentation was counterbalanced across participants. Participants were instructed to either respond as quickly as possible to the response cue or to withhold a response, and that they should explore different actions and learn which response was correct, given the previous cue shape. The tasks were presented and responses collected using Presentation software. The task paused every 60-90 secs to avoid fatigue and allow the participant to blink and move their eyes.

To measure Pavlovian biases across the different conditions within a task, and compare across tasks, we calculated the following measures using the formulae described by Cavanagh and colleagues (2013): Reward Invigoration (RI) was calculated as (Go on Go to Win + No-Go to Win) / Total Go; Punishment Suppression (PS) calculated as (No-Go on Go to Avoid + No-Go to Avoid) / Total No-Go; and Pavlovian Performance Bias (PPB) as the average of RI and PS. For each of these measures, a bias score of 50% indicates no Pavlovian bias and performance based only on an instrumental system, whereas a bias score greater than 50% indicates the presence of Pavlovian bias.

Bayesian statistical analysis of behavioral data was performed using the Stan probabilistic programming language (Carpenter et al., 2016) and the MatlabStan interface (Lau, 2014). Accuracy on the Go/NoGo task was analyzed using a linear mixed effects model with varying intercept and slopes for action (Go, NoGo), conflict (Conflict, No Conflict), and outcome (wins, losses). Robust Pearson's correlations with uncertainty in measurement were estimated using a modification of procedure described by Lee and Wagenmakers (2013). The No-U-Turn (NUTS) sampler, a variant of Hamiltonian Monte Carlo (HMC) sampling implemented in Stan was used to estimate posterior distributions. We ran two chains of 2000 samples each, discarding the first 1000 samples as burn-in. Priors were weakly informative: Gaussian *N*(0,100) for means, *Cauchy* (0, 5) for standard deviation (Gelman et al., 2013). For the robust Pearson's correlation, a multivariate *Student's t* prior was used with an *Exponential(1/29)* hyperprior for the degrees of freedom. Inferences were based on 95% highest density intervals (HDIs) of posterior distributions.

### EEG Recording and Analysis

High-density EEG was recorded from 122 scalp channels and horizontal and vertical electrooculogram (EOG) channels relative to a vertex reference using an electrocap (Neuroscan; Compumedics, Abbotsford, Australia). A Synamps2 system (Compumedics) was used to record data at a sampling rate of 1000 Hz with a bandpass filter of 0.1–100 Hz. Following data acquisition, the locations of each electrode and the three major fiducial points (left and right periauricular points and nasion) were digitized using a Polaris optical camera (Northern Digital, Waterloo, Canada) with Brainsight software (Rogue Research, Montreal, Canada). Importantly, during digitization a reference was attached to the participant's head so that head movements would not result in inconsistent measurements.

Data were then analyzed using EEGLAB, FieldTrip, and custom Matlab functions. Data were first epoched around the onset of the shape cue (-1000ms to +1500ms), inspected for bad channels (range: 0-4 electrodes rejected and interpolated) and epochs with gross artifact to be rejected (range: 0-6 epochs). Data were referenced to the common average, zero-meaned, and low pass filtered at 40Hz. Independent Components Analysis (ICA), using the Informax algorithm implemented in EEGLAB was then run on the 122 scalp electrodes to identify eye blinks and movements and muscle artifacts. Semi-Automatic Selection of Independent Components of the electroencephalogram for Artifact correction (SASICA) (Chaumon et al., 2015) was used to aid the identification of artifact components. Following ICA, data were resampled to 250Hz. Time frequency analysis of EEG data from -1000ms to +1500ms relative to the onset of the shape cue for each condition from three regions of interest (frontocentral, left posterior, right posterior) was performed in FieldTrip using Morlet wavelets (length = 5 cycles), with frequencies between 4 and 30Hz in 0.5Hz steps. Trials were averaged and decibel (10*log10) power relative to a baseline from -500ms to -200ms before the shape cue onset was calculated. Band-limited power in theta (4-7Hz), alpha (8-14Hz), and beta (15-30Hz) bands for each trial was calculated using zero-phase, two-pass Butterworth IIR filters, followed by Hilbert transformation. Trials were averaged and power in each frequency band was then converted to percent change from baseline using the baseline period of -500ms to -200ms before the onset of the shape cue.

Analyses of the effects of Pavlovian-Instrumental conflict on EEG power were conducted by averaging the two conflict conditions (Go to Avoid Loss, No-Go to Win) and the two no conflict conditions (Go to Win, No-Go to Avoid Loss), and comparing the two conditions using t-tests at each time point from the onset of the shape cue to +600ms after onset, at each electrode and each frequency band. We then examined the relationship between Pavlovian bias (PPB and model parameters) using Spearman's rank ordered correlation at each time point and electrode, separately for theta, alpha, and beta frequency bands. We used cluster-based permutation testing to correct for multiple comparisons, with 2000 permutations, a cluster forming threshold of p<0.05, and a significance threshold of p<0.05. For computational modeling, single trial estimates of shape-cue related EEG power in each band relative to the pretrial baseline were extracted from the three regions of interest for each condition. Single trial data were averaged within early (100-300ms) and later (300-500ms) windows. These data were then z-normalized prior to inclusion in the computational models.

### Computational Modeling

We used Bayesian hierarchical modeling and stepwise model comparison to examine the mechanism by which Pavlovian bias influences choice behavior in the Social Go/No-Go and Monetary Go/No-Go tasks, and to determine the nature of the relations between EEG in the theta and alpha bands and Pavlovian bias. Separate RL models were fit for the Social Go/No-Go and Monetary Go/No-Go tasks. We adapted a series of existing computational models of this task (Guitart-Masip et al., 2012; Cavanagh et al., 2013) to estimate parameters that underlie Pavlovian-Instrumental interactions during social and monetary RL. Progressively more complex models were fit to trial-by-trial behavioral data using hierarchical Bayesian model fitting implemented in Stan (Carpenter et al., 2016), and models were compared by examining if they explained additional variance (penalized for additional parameters).

Performance on the social and monetary Go/No-Go tasks was first modeled as an Instrumental (Q) learning process with state-action (Q) values updated on a trial-by-trial basis using a Q-learning algorithm [44]:

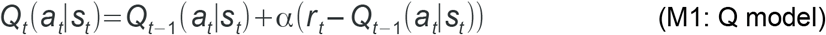

where reinforcements (*r*) come from the set *r*∈(−1,01) and a is a free parameter accounting for learning rate, bound between 0 and 1. To account for bias to Go, regardless of the task conditions, a second model included an additional free parameter *(b)* on Go trials:

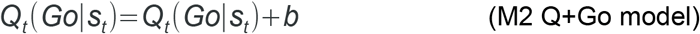

To model the influence of Pavlovian bias, we modeled the learned value of each stimulus as a function of reward history:

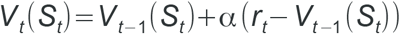

The value of each stimulus was then added to bias the state-action value for Go responses:

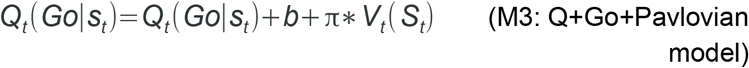

where π is free parameter representing the strength of Pavlovian bias.

As in Cavanagh and colleagues (2013), we examined if frontal EEG oscillatory power reflected the suppression of Pavlovian influences by comparing an additional set of three models. Each of these models included an effect of EEG power free parameter (ρ), that weighted the contribution of trial-by-trial EEG power (theta, alpha) to the balance between the Instrumental controller (Q) and the Pavlovian controller (*V*). As the influence of Pavlovian bias was expected only on trials in which Pavlovian controller conflicted with the Instrumental controller, the contribution of EEG power was only modeled for conflict trials. The first model examined if EEG power directly modulates Pavlovian influence:

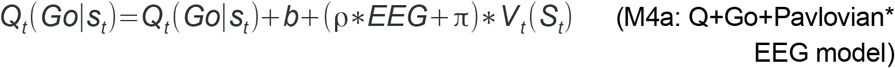

The second model examined if EEG power modulates the influence of the Instrumental (Q) system:

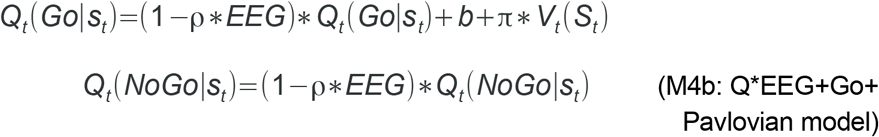

A third model examined if EEG modulates a trade-off between Pavlovian and Instrumental systems:

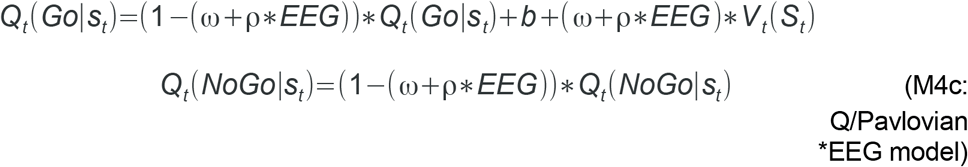

The *Q* values were used to control the choice probability of action according to a softmax rule:

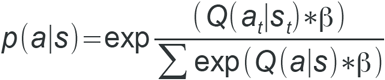

where β is an “inverse temperature” parameter that reflects the degree to which the probability reflects the highest value action. Bayesian hierarchical modeling was used to estimate the joint posterior distribution of parameters for the Q model (α, β), the Q+Go model (α, β *,b*) the Q+Go+Pavlovian bias model (α, β, *b,* π), and the three models that included trial-by-trial EEG (Q+Go+Pavlovian*EEG; Q*EEG+Go+Pavlovian;Q/Pavlovian*EEG), conditional on all participant choices and rewards. The No-U-Turn (NUTS) sampler implemented in Stan was used to estimate posterior distributions of parameters. We ran two chains of 2000 samples each, discarding the first 1000 samples as burn-in. All of the free parameters were treated as random effects. Models were compared using the Widely-Applicable Information Criterion (WAIC; Watanabe, 2013). The different behavior-only computational models were compared to the simplest behavioral model, while different behavior + EEG computational models were compared to the best fitting behavior-only model. Posterior predictive model checking was performed by generating one step ahead predictions of action probabilities for trial n, based on model parameters and choice/outcome relations for trials 1 to (n-1). Action probabilities for each trial for each of the 2000 samples were averaged for each participant. Separate posterior distributions of parameters were modeled for the social and monetary tasks. Weakly informative priors were used: priors for parameters with infinite support (β, *b,* π, ω, ρ) were Gaussian N(0,100) for means, *Cauchy* (0, 5) for standard deviation; for parameters bound between [0,1] (α), priors were drawn from *Beta* (A,B) distributions, with A and B transformed from M=A/(A+B) and S= 1/sqrt(A+B), with M drawn from a uniform distribution *U* (0,1) and S drawn from uniform distribution U (0, Inf).

## Results

The probability of making a Go response was calculated for each condition, separately for each task. Plots of the average p(Go) across trials indicate that participants were more likely to ‘go’ to the response cue when this response was associated with reward (GTW), and were less likely to ‘go’ when the response was associated with a loss (NGTA) (Figure 2A). This pattern was consistent across the Social and Monetary tasks. To facilitate a direct comparison of performance across conditions, we used p(Go) as a measure of correct response for the Go trials, and 1-p(Go) for correct response for the No-Go trials (Figure 2B). Across both tasks, accuracy performance was consistent with an influence of Pavlovian bias. Accuracy was higher in the conditions in which there was no Pavlovian-Instrumental conflict (Go to Win, No-Go to Avoid) than in the conditions in which there was conflict (Go to Avoid, No-Go to Win), for both the Social task (posterior mean slope = 10.72 [4.17, 17.52]) and the Monetary task (posterior mean slope = 11.33 [0.11, 21.93]).

**Figure 2.**
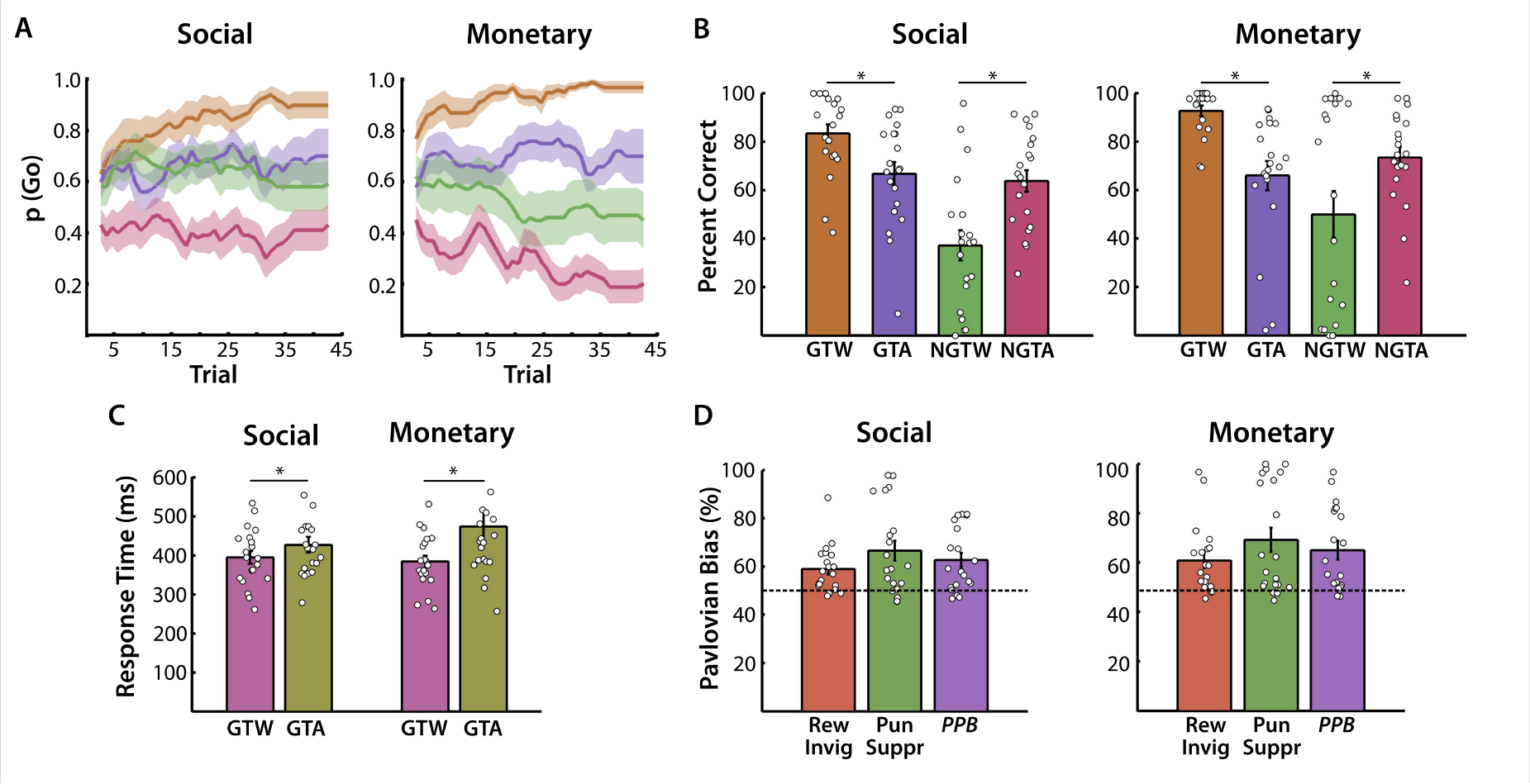
Behavioral results from the Social and Monetary Go/NoGo tasks. **A.** Trial-by-trial average (+/- s.e.) probability of making a ‘go’ response to the response cue (5 trial moving average). Burnt orange is Go to Win (GTW), dark lilac is Go to Avoid (GTA), mantis green is NoGo to Win (NGTW), and dark terra cotta NoGo to Avoid (NGTA). **B.** Average (+/- s.e.) and individual participant percent correct responses (p(Go) for Go trials, 1-p(Go) for NoGo trials). **C.** Average (+/- s.e.) and individual participant reaction times for ‘go’ responses for GTW and GTA trials. **D.** Average (+/- s.e.) and individual participant measures of Pavlovian bias (Rew Invig = reward invigoration; Pun Suppr = Punishment Suppression; PPB = Pavlovian Performance Bias). Colorblind friendly palettes were created using IwantHue (http://tools.medialab.sciences-po.fr/iwanthue/) and k-means clustering.

Accuracy was higher in the Go trials than in the No-Go trials for the Social task (posterior mean slope = 12.46 [2.5, 19.00]), but not the Monetary task (posterior mean slope = 9.07 [−2.36, 20.64]). Overall, accuracy was not higher for wins than in losses for either the Social task (posterior mean slope = −2.50 [−7.68, 3.04]) and the Monetary task (posterior mean slope = 1.13 [-7.48, 11.19]). In further support of the presence of Pavlovian bias influencing performance, we observed that (log transformed) reaction times (RT) for Go trials with no conflict (Go to Win) were faster than Go trials with conflict (Go to Avoid) for both the Social task (posterior mean difference = −0.08 [−0.14, −0.01]) and the Monetary task (posterior mean difference = −0.16 [−0.31, −0.02]) (Figure 2C).

To quantify Pavlovian influence on choice on the two tasks, we calculated indices of Reward Invigoration (RI), Punishment Suppression (PS), and overall Pavlovian Performance Bias (PPB). For each of these measures, a bias score of 50% indicates no Pavlovian bias and performance based only on an instrumental system, whereas a bias score greater than 50% indicates the presence of Pavlovian bias. The posterior 95% HDIs of the means for the three indices were greater than 50% for the Social task (RI = [54.38, 62.67]; PS = [57.74, 74.45]; PPB = [56.68, 68.15]) and the Monetary task (RI = [54.26, 66.91]; PS = [57.45, 78.31]; PPB = [56.59, 76.02]) (Figure 2D). The pattern of accuracy and PPB observed on the Social and Monetary Go/No-Go tasks used in the present study were qualitatively similar to previous studies employing versions of the Monetary task (Cavanagh et al., 2013; Guitart-Masip et al., 2012). As was the case with these studies, there was considerable variability in the extent to which Pavlovian biases influenced performance, especially in the No-Go to Win conditions (Figure 2B). We examined if the Pavlovian bias observed on both the Social and Monetary tasks reflected a common learning mechanism, regardless of the properties of the outcome, by examining if individual differences in Pavlovian Bias across was correlated across Social and Monetary tasks. Results indicated that RI (posterior mean r = 0.24 [-0.49, 0.87]), PS (posterior mean r = 0.08 [-0.91, 0.93]), and PPB (posterior mean r = 0.13 [-0.79, 0.88]) were not reliably correlated across the Social and Monetary tasks, indicating that those participants who were most influenced by social outcomes were not similarly affected by monetary outcomes (and vice versa).

### EEG

We first examined if Pavlovian-Instrumental conflict during Social or Monetary reinforcement learning was associated with changes to EEG power by comparing conditions with Conflict (GTA and NGTW) to conditions with No Conflict (GTW and NGTA), averaged across all trials in each condition. We examined EEG power locked to the onset of the abstract shapes that had no initial value prior to the start of the task (Figure 3A). It should be noted that the association with a motor response and the valence of outcome was balanced across Conflict and No Conflict conditions. No significant differences in EEG power between Conflict and No Conflict conditions were observed in either the Social Go/No-Go task or the Monetary Go/No-Go task. To examine the relationship between Pavlovian biases in behavior and conflict-related changes in EEG power, we correlated behavioral performance with conflict-specific (Conflict - No Conflict) EEG responses in the theta, alpha, and beta frequency bands for the Social and Monetary Go/No-Go tasks.

**Figure 3.**
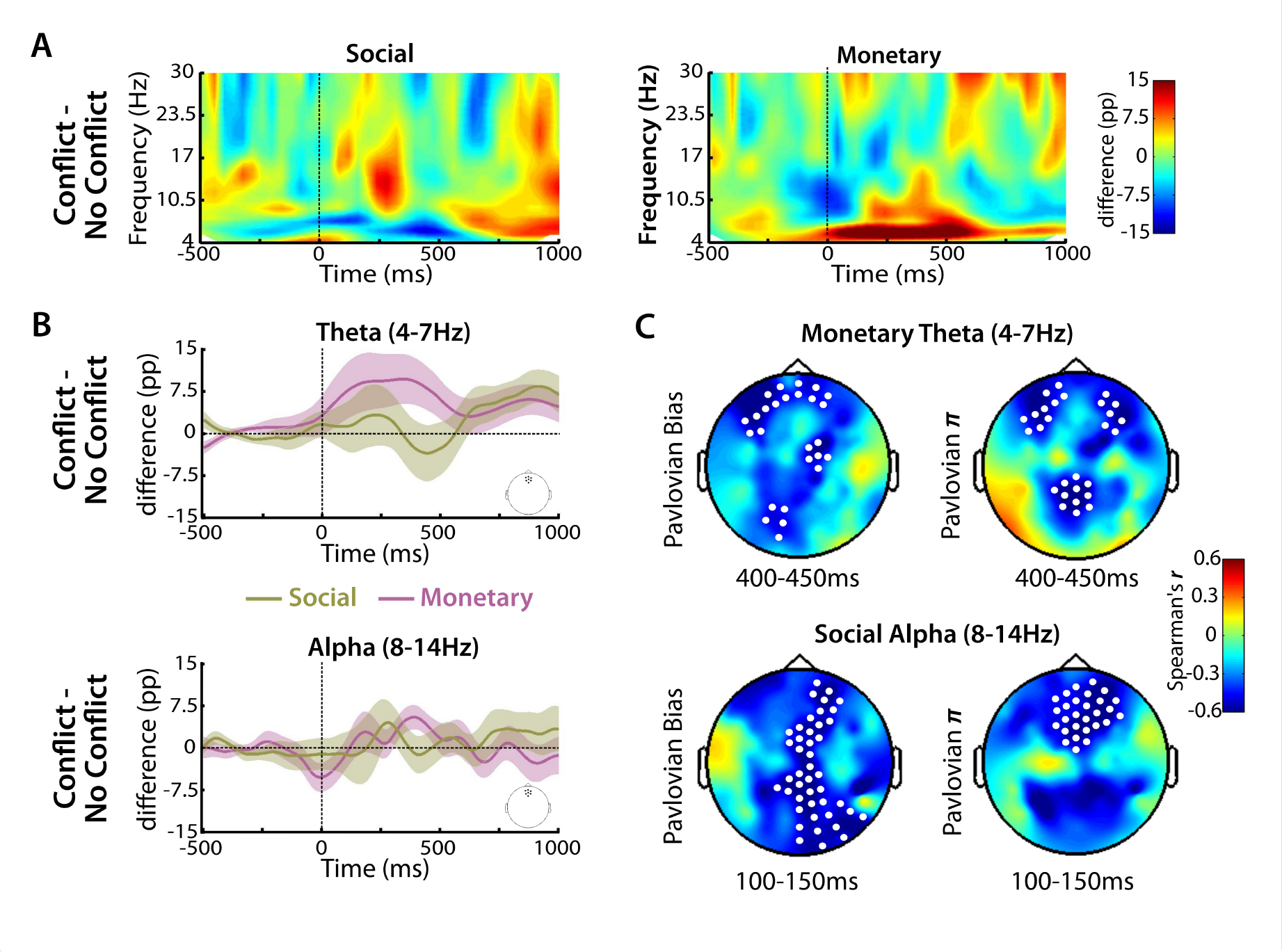
EEG Data Time Locked to Onset of the Shape Cue. A. Time-frequency plots of the average difference between Pavlovian Conflict (GTA, NGTW) and No Conflict (GTW, NGTA) from frontocentral ROI (pp = percentage points). B. Average (+/-) difference in bandpass filtered power in theta and alpha bands between Conflict and No Conflict conditions from frontocentral ROI. C. Maps of the correlation (Spearman's r) between Pavlovian bias and EEG power (Conflict - No Conflict). Top row shows correlation maps (and electrode sites within significant clusters) between Pavlovian Performance Bias and theta power (left), and model-based Pavlovian bias parameter π and theta power (right), from the Monetary task. Bottom row shows correlation maps (and electrode sites within significant clusters) between Pavlovian Performance Bias and alpha power (left), and model-based Pavlovian bias parameter π and alpha power (right), from the Social task.

A previous study reported significant negative correlation between conflict-specific frontocentral theta EEG power and the behavioral PPB measure from a monetary Go/No-Go task (Cavanagh et al., 2013). We found significant negative correlation between conflict-specific theta EEG power and the PPB measure in the Monetary Go/No-Go task in frontal and parietal sites (cluster: 200-528ms; p=0.002 corrected) (Figure 3C). This is later than the timing of the correlation reported by Cavanagh and colleagues (2003) of 175-350ms, but consistent with the timing of conflict-related activity in the theta band reported elsewhere (Cohen & Donner, 2013). This correlation was restricted to the theta band and no significant correlation between PPB and alpha power was observed for the Monetary Go/No-Go task. In contrast to the findings from the Monetary Go/No-Go task, no significant correlation between PPB and conflict-specific theta was observed for the Social Go/No-Go task. Instead, a significant negative correlation with PPB was observed in the alpha band over frontal and posterior sites (cluster: 20-268ms; p=0.005 corrected) (Figure 3C). No significant correlations between PPB from either the Social Go/No-Go task or the Monetary Go/No-Go task were observed in the beta frequency band.

### Computational Modeling

Beginning first with behavior-only RL models, we observed that relative to a Q-learning (M1) and the Q-learning+Go (M2) models, including a Pavlovian bias parameter (π) increased model fit (i.e. decreased WAIC relative to the Q model) of M3 considerably for both the Social Go/No-Go task and the Monetary Go/No-Go task (Table 1). To examine the relationship between model estimates of Pavlovian bias and EEG activity, we correlated the model parameter π and EEG power in the theta and alpha frequency bands for the Social Go/No-Go and Monetary Go/No-Go tasks. We observed significant negative correlation between π and alpha power for the Social Go/No-Go task over frontal sites (cluster: 64-280ms; p=0.007 corrected), and between π and theta power for the Monetary Go/No-Go task over frontal sites (cluster: 200-528ms; p=0.02 corrected) (Figure 3C). These effects indicate that individuals with greater level of theta power (for Monetary Go/No-Go) or alpha power (for Social Go/No-Go) show less Pavlovian bias during learning, however these results do not reveal the mechanism underlying trial-by-trial interactions between Pavlovian and Instrumental systems. To examine trial-by-trial effects, we compared three models in which EEG power in either the theta or alpha band modulated the Pavlovian influence directly (M4a: Q+Go+Pavlovian*EEG), modulated the Instrumental influence (M4b: Q*EEG+Go+Pavlovian), or modulated the trade-off between Pavlovian and Instrumental influences (M4c: Q/Pavlovian*EEG + Go).

**Table 1.** Widely Applicable Information Criterion (WAIC) for the different computational models from the Social and Monetary tasks.

|  | **Social** | **Monetary** |
| --- | --- | --- |
| **Model** | **WAIC** | **WAIC** |
| **M1:** Q | 4092 | 3525 |
| **M2:** Q + Go | 3661 | 3014 |
| **M3:** Q + Go + Pav | 3373 | 2747 |
| **M4a:** Q + Go + Pav * EEG (alpha) | **3358** | 2714 |
| **M4a:** Q + Go + Pav * EEG (theta) | 3371 | **2697** |
| **M4b:** Q * EEG + Go + Pav (alpha) | 3364 |  |
| **M4c:** Q/Pav * EEG + Go (alpha) | 3366 |  |
| **M4b:** Q * EEG + Go + Pav (theta) |  | 2938 |
| **M4c:** Q/Pav * EEG + Go (theta) |  | 2747 |

Results indicated that for the Social Go/No-Go task, the winning model (M4a) was one in which the influence of the Pavlovian system was modulated on a trial-by-trial basis by frontal alpha EEG power from a 100-300ms window after shape onset (Table 1). For the Monetary Go/No-Go task, results indicated that instead the winning model (M4a) was one in which the influence of the Pavlovian system was modulated on a trial-by-trial basis by frontal theta EEG power in a 300- 500ms window after shape onset (Table 1).

These effects showed some level of frequency-dependency, as models that used theta EEG power to modulate Pavlovian bias were a poor fit for data from the Social Go/No-Go task, whereas models using alpha EEG power to modulate Pavlovian bias were a poor fit for data from the Monetary Go/No-Go task. Posterior distributions (including posterior mean and 95% Highest Density Intervals) of the group-level parameters, and plots of posterior means of individual-level parameter estimates, of the winning model for the Social and Monetary Go/No-Go tasks are presented in Figure 4A.

**Figure 4.**
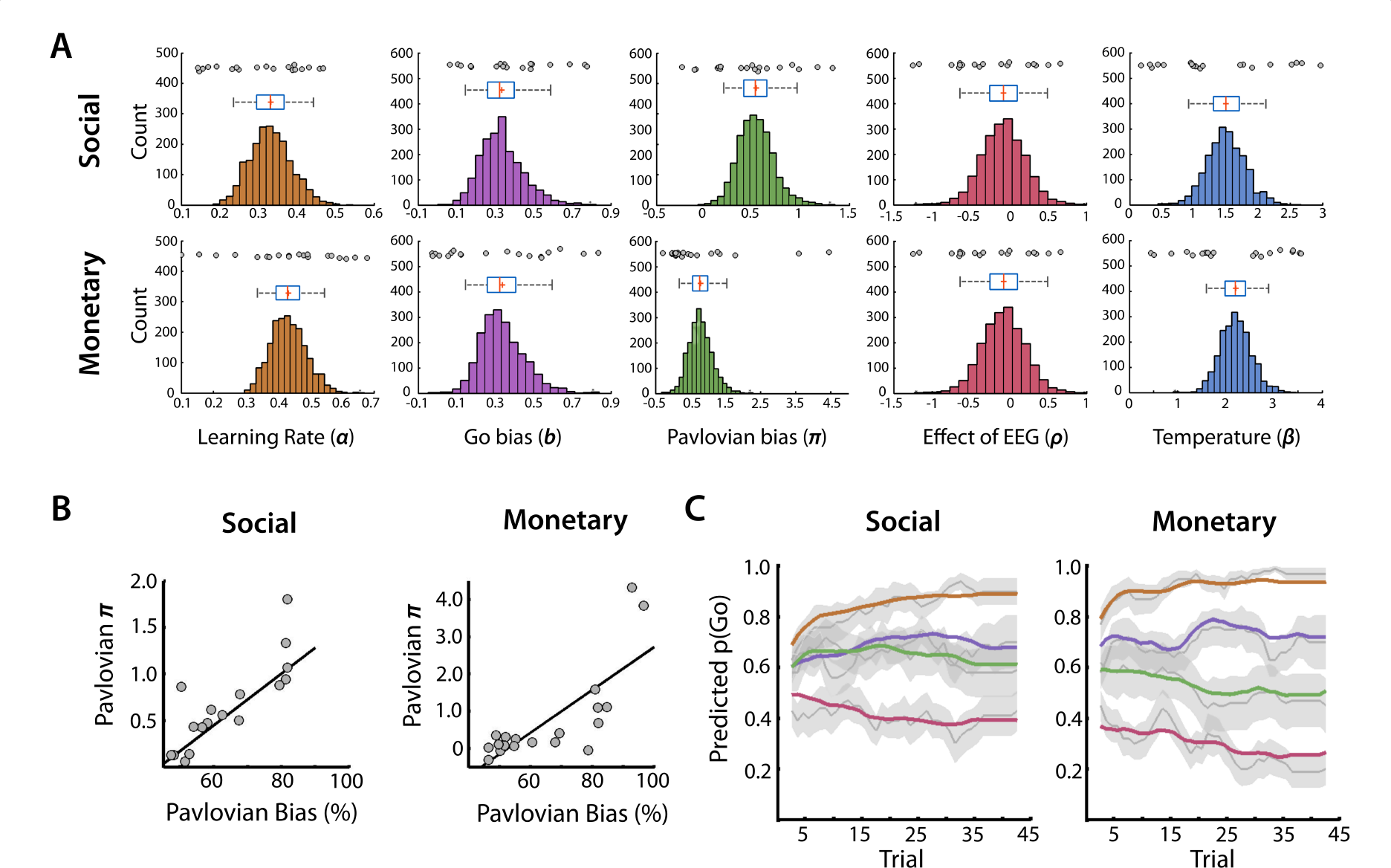
Posterior Estimates of Model Parameters and Posterior Predictive Checks. A. Posterior distributions of model parameters from the M4a model from the Social (top row) and Monetary (bottom row) tasks. Histograms and box and whisker plots represent posterior distribution of HMC samples (n = 2000) of the group-level parameters. Box indicates posterior interquartile range (red line = median, red cross = mean), whiskers indicate 95% HDI. Circles indicate posterior means for participant-level parameters for each participant. B. Scatterplots and least squares line of Pavlovian Performance Bias (x axis) and model-based Pavlovian parameter π (y axis) from Social (left) and Monetary (right) tasks. C. Average trial-by-trial one step ahead predictions (5 trial moving average) of the probability of a ‘go’ response from the four conditions of the Social (left) and Monetary (right) tasks. Colors of conditions are the same as in Figure 2A. Grey lines represent average (+/- s.e) of participants' probability of ‘go’ response.

As a posterior predictive check, the posterior mean of the Pavlovian model parameter n was calculated for each participant and correlated with PPB for the Social Go/No-Go task and the Monetary Go/No-Go task (Figure 4B). Results indicated that the model parameter n was correlated with PPB for both the Social Go/No-Go (posterior mean r = 0.77 [0.40, 0.98]) and Monetary Go/No-Go (posterior mean r = 0. 74 [0.45, 0.96]) tasks. There were two outliers in the distribution of π from the Monetary Go/No-Go task; while the robust Pearson correlation is generally robust to such outliers, we repeated the correlation after removing the two outliers and found only minimal change to the correlation (posterior mean r = 0.73 [.37, 0.99]). We also generated one step ahead predictions of choice for trial tn for each participant, from the trial-by-trial combination at trials t1 to tn-1 of model estimates, behavioral choice/outcomes, and EEG data (Figure 4C). These predictions capture key features of the behavioral data, including an increased probability to Go over NoGo responses, and Pavlovian bias, for both the Social Go/NoGo and the Monetary Go/NoGo tasks.

## Discussion

Nonverbal social signals can shape our behavior by providing a signal of the value of our actions to another: a friendly smile can encourage us to continue, while an angry scowl can stop us in our tracks (Fridja, 1986; Van Kleef, 2009). Expressions of emotions in other people, conveyed through facial expressions, can also generate affective responses and elicit associated automatic approach or avoidance behaviors from an observer (Marsh et al., 2005; Seidel et al., 2010). It can therefore be useful to learn what cues predict the expression of emotions in others. At times, the automatic responses to reward or punishment (and/or the cues that predict these outcomes), and the instrumental demands of a social interaction, can come into conflict. In this study, we adapted a computational model that proposes distinct motivation systems for action-value (Instrumental) learning and Pavlovian learning of stimulus-outcome associations (Dayan et al, 2006), to examine reinforcement learning from social feedback from others. We found that conflict between Pavlovian and Instrumental demands can impair learning accuracy and response times when learning cue-action relations based on social feedback. Moreover, the adverse effects of Pavlovian-Instrumental conflict were due to the trial-by-trial contribution of Pavlovian biases to approach social reward or withdrawal from social punishment. The interactions between Pavlovian and Instrumental systems during social reinforcement learning were computationally very similar to those observed during reinforcement learning from monetary outcomes, although we observed that across participants the Pavlovian bias for social outcomes was not correlated with Pavlovian bias for monetary outcomes. We also observed distinct brain oscillatory mechanisms underlying the control of social and monetary outcome biases.

Controlling responses to social feedback is important for adapting behavior to different social contexts. While feedback from others can come in the form of verbal or physical cues, signaling the value of behavior through an emotional facial expression can be a quick, and often effective, means of communicating whether a behavior is socially appropriate or not (Niedenthal & Brauer, 2012; Van Kleef, 2009). Defining the control of socially-based emotions in computational terms is important for a detailed understanding of this phenomenon, including understanding individual differences and the circumstances in which emotion control breaks down (Etkin et al., 2016; Raio et al., 2016). The interaction between Pavlovian and Instrumental motivation systems described in the present study represents only one way in which motivation systems might coordinate and control behavior. The computational approach we used represents both Pavlovian and Instrumental systems each as model-free, assigning values to stimuli and actions based on the outcome history rather than any explicit, causal representation of the world (Doll et al., 2012; O’Doherty et al., 2017). This is likely an oversimplification, as model-based strategies play an important role in controlling behavior including learning from social feedback (Doll et al., 2012; Dayan & Berridge, 2014). Interestingly, a recent study indicated that the strength of model-based learning in a reinforcement learning task was negatively correlated with the strength of Pavlovian to Instrumental transfer, in tasks using monetary outcomes (Sebold et al., 2016). These findings suggest that model-based Instrumental learning and the control of Pavlovian biases might share a common basis.

We observed that successful neural control of the Pavlovian system during learning from social feedback was associated with increased oscillatory EEG activity in the alpha (8-14Hz) frequency range, particularly over frontal sites. A brief increase in frontal alpha EEG power in the first 100-300ms after the onset of a predictive cue was associated on a trial-by-trial basis with the suppression of Pavlovian bias, and, across participants, the extent of Pavlovian influence on behavior was associated with increases in alpha power to cues associated with P-I conflict in this same time window. This study is the first to describe in computational terms the neural basis of the control of Pavlovian bias during Instrumental learning from social feedback. We also observed that learning from monetary feedback involved similar computational processes to learning from social feedback, but theta (4-7Hz) activity was involved for the control of Pavlovian bias to monetary feedback. Successful suppression of Pavlovian bias was associated with an increase in frontal theta EEG activity in a time window later than that for the social task. The results from the monetary task are largely consistent with a previous report using a very similar task by Cavanagh and colleagues (2013), although these authors found frontal theta-based suppression of Pavlovian bias following monetary feedback in an earlier time window (175-350ms) to that observed for the monetary feedback in the present study.

Dynamic oscillatory EEG and MEG responses likely reflect the integration of local circuit activity with large-scale coordinated activity across networks of regions, rather than the response of any one brain region alone (Freeman, 1975; Fries, 2015; Nunez, 2000). We observed that midfrontal alpha and theta played very similar roles in the suppression of Pavlovian bias during social and monetary reinforcement learning, respectively, albeit with distinct time courses. Two explanations for these findings seem likely: first, the frequency-dependent effects we observed could reflect two quite distinct neural networks. Midfrontal theta has long been associated with the engagement of cognitive control, and appears to reflect the activity of a network included medial PFC, dorsal anterior cingulate cortex, and basal ganglia (Cavanagh & Frank, 2013). The sources and role of alpha oscillations have been subject to considerably more debate. One hypothesis is that alpha power reflects the engagement of active inhibition of task or goal-irrelevant representations (Klimesch et al., 2007; Jensen and Mazaheri, 2010; Bollimunta et al., 2011; Sadaghani & Kleinschmidt, 2016). For example, the pulsed inhibition theory of cortical alpha proposes that pulses of top-down inhibition at approximately every 100ms act to suppress task-irrelevant representations in sensory and motor cortex (Klimesch et al., 2007; Haegens et al., 2014). It is possible that the network associated with suppression of biases to approach social reward-predicting stimuli and withdrawal from social punishment-predicting stimuli during learning is engaged early and consists of alpha-frequency pulses of inhibition from frontal regions to sensorimotor representations, distinct from a frontostriatal network involved in suppression of Pavlovian bias associated with monetary outcomes.

An alternative explanation for the present findings is that a largely overlapping network is recruited for the suppression of Pavlovian biases associated with social and monetary outcomes. This network might recruit distinct subregions for the control of behaviors associated with different classes of outcomes, with brain regions preferentially involved in the determining the social importance of environmental cues, including in medial prefrontal and lateral temporal cortices, engaged during the social task. Recruitment of distinct brain regions into a network might be expected to shift the dominant oscillatory rhythm of that network, either because of the intrinsic temporal properties of those regions or through reconfiguration of longer-range connections throughout the network (including shortening or lengthening of paths) (Freeman, 1975; Nunez, 2000). In addition, the early timing of the alpha frequency-based Pavlovian suppression mechanism could reflect a greater priority given for primary (e.g., social) reinforcenrs relative to secondary reinforcers (e.g., money) (Anderson, 2016; Kringelbach & Rolls, 2003).

Together, our findings suggest a computational mechanism though which the emotional responses to social signals from other people are balanced with the instrumental value that those signals provide. While this mechanism shares similarities with that used to suppress biases associated with other forms of reward, the control of Pavlovian bias from social outcomes was associated with a distinct basis in brain oscillatory activity. These results provide details of how controlling how we feel in response to the feedback from others can be important for learning behaviors appropriate for social contexts, and show how tracking the dynamics of brain activity, combined with computational modeling, can reveal how such control is implemented.

## References

Anderson BA. (2016). Social reward shapes attentional biases. Cogn Neurosci. 7:30–6.

Apps MA, Rushworth MF, Chang SW. (2016). The Anterior Cingulate Gyrus and Social Cognition: Tracking the Motivation of Others. Neuron. 90:692–707.

Behrens TE, Hunt LT, Rushworth MF. (2009). The computation of social behavior. Science. 324:1160–4.

Berridge KC. (1996). Food reward: brain substrates of wanting and liking. Neurosci Biobehav Rev. 20:1–25.

Bollimunta A, Mo J, Schroeder CE, Ding M. (2011) Neuronal mechanisms and attentional modulation of corticothalamic α oscillations. J Neurosci. 31:4935–43.

Bray S, Rangel A, Shimojo S, Balleine B, O’Doherty JP. (2008). The neural mechanisms underlying the influence of pavlovian cues on human decision making. J Neurosci. 28:5861–6.

Buschman TJ, Miller EK. (2014). Goal-direction and top-down control. Philos Trans R Soc Lond B Biol Sci. 369: 20130471.

Carpenter, B, Gelman, A, Hoffman, M, Lee, D, Goodrich, B, Betancourt, M, Brubaker, MA, Guo, J, Li, P, Riddell, A. (2017). Stan: A probabilistic programming language. Journal of Statistical Software, 76 (1):1–32.

Cavanagh, JF, Eisenberg, I, Guitart-Masip, M, Huys, Q, Frank, MJ. (2013). Frontal theta overrides pavlovian learning biases. Journal of Neuroscience. 33(19):8541–8

Cavanagh JF, Frank MJ. (2014) Frontal theta as a mechanism for cognitive control. Trends Cogn Sci. 18:414–21.

Chaumon M, Bishop DV, Busch NA. (2015). A practical guide to the selection of independent components of the electroencephalogram for artifact correction. J Neurosci Methods. 250:47–63.

Cohen MX, Donner TH. (2013). Midfrontal conflict-related theta-band power reflects neural oscillations that predict behavior. J Neurophysiol. 110:2752–63.

Dayan P, Berridge KC. (2014). Model-based and model-free Pavlovian reward learning: revaluation, revision, and revelation. Cogn Affect Behav Neurosci. 14:473–92.

Dayan, P, Niv, Y, Seymour, B, Daw, ND. (2006). The misbehavior of value and the discipline of the will. Neural Networks. 19(8):1153–60.

Doll BB, Simon DA, Daw ND. (2012). The ubiquity of model-based reinforcement learning. Curr Opin Neurobiol. 22:1075–81.

Etkin A, Büchel C, Gross JJ. (2015). The neural bases of emotion regulation. Nat Rev Neurosci. 16:693–700.

Etkin A, Egner T, Peraza DM, Kandel ER, Hirsch J. (2006). Resolving emotional conflict: a role for the rostral anterior cingulate cortex in modulating activity in the amygdala. Neuron. 51:871–82.

Fries P. (2015). Rhythms for Cognition: Communication through Coherence. Neuron. 88:220–35.

Frijda, NH. (1986). The emotions. Cambridge, UK: Cambridge University Press.

Gelman A, Carlin JB, Stern HS, Dunson DB, Vehtari A, Rubin DB (2013). Bayesian Data Analysis, Third Edition. Chapman Hall, Boca Raton, FL.

Geurts DE, Huys QJ, den Ouden HE, Cools R. (2013). Aversive Pavlovian control of instrumental behavior in humans. J Cogn Neurosci. 25:1428–41.

Guitart-Masip, M, Huys, QJ, Fuentemilla, L, Dayan, P, Duzel, E, Dolan, RJ. (2012). Go and no-go learning in reward and punishment: interactions between affect and effect. Neuroimage. 62(1):154–66.

Haegens S, Vázquez Y, Zainos A, Alvarez M, Jensen O, Romo R. (2014). Thalamocortical rhythms during a vibrotactile detection task. Proc Natl Acad Sci U S A. 111:E1797–805.

Helfrich RF, Knight RT. (2016). Oscillatory Dynamics of Prefrontal Cognitive Control. Trends Cogn Sci. 20:916–930.

Huys QJ, Cools R, Gölzer M, Friedel E, Heinz A, Dolan RJ, Dayan P. (2011). Disentangling the roles of approach, activation and valence in instrumental and pavlovian responding. PLoS Comput Biol. 7:e1002028.

Hwang K, Ghuman AS, Manoach DS, Jones SR, Luna B. (2014). Cortical neurodynamics of inhibitory control. J Neurosci. 34:9551–61.

Jensen O, Mazaheri A. (2010). Shaping functional architecture by oscillatory alpha activity: gating by inhibition. Front Hum Neurosci. 4:186.

Klimesch W, Sauseng P, Hanslmayr S, Gruber W, Freunberger R. (2007). Event-related phase reorganization may explain evoked neural dynamics. Neurosci Biobehav Rev. 31:1003–16.

Kringelbach ML, Rolls ET. (2003). Neural correlates of rapid reversal learning in a simple model of human social interaction. Neuroimage. 20:1371–83.

Lee and Wagenmakers (2013) Bayesian Cognitive Modeling: A Practical Course Lee MD, Wagenmakers E-J. Cambridge: Cambridge University Press

Lin A, Adolphs R, Rangel A. (2012). Social and monetary reward learning engage overlapping neural substrates. Soc Cogn Affect Neurosci. 7:274–81.

Mackintosh, NJ. (1983). Conditioning and Associative Learning. Oxford, UK: Oxford University Press.

Marsh AA, Ambady N, Kleck RE. (2005). The effects of fear and anger facial expressions on approach and avoidance-related behaviors. Emotion. 5:119–24.

Niedenthal PM, Brauer M. (2012). Social functionality of human emotion. Annu Rev Psychol. 2012;63:259–85.

Nunez PL, Wingeier BM, Silberstein RB. (2001). Spatial-temporal structures of human alpha rhythms: theory, microcurrent sources, multiscale measurements, and global binding of local networks. Hum Brain Mapp. 13:125–64.

O’Doherty JP, Cockburn J, Pauli WM. (2017). Learning, Reward, and Decision Making. Annu Rev Psychol. 68:73–100.

Prévost C, Liljeholm M, Tyszka JM, O’Doherty JP. (2012). Neural correlates of specific and general Pavlovian-to-Instrumental Transfer within human amygdalar subregions: a high-resolution fMRI study. J Neurosci. 32:8383–90.

Raio CM, Goldfarb EV, Lempert KM, Sokol-Hessner P. (2016). Classifying emotion regulation strategies. Nat Rev Neurosci. 17:532.

Rescorla RA, Solomon RL. (1967). Two-process learning theory: Relationships between Pavlovian conditioning and instrumental learning. Psychol Rev. 74:151–82.

Sadaghiani S, Kleinschmidt A. (2016). Brain Networks and α-Oscillations: Structural and Functional Foundations of Cognitive Control. Trends Cogn Sci. 20:805–817.

Sebold M, Schad DJ, Nebe S, Garbusow M, Jünger E, Kroemer NB, Kathmann N, Zimmermann US, Smolka MN, Rapp MA, Heinz A, Huys QJ. (2016). Don’t Think, Just Feel the Music: Individuals with Strong Pavlovian-to-Instrumental Transfer Effects Rely Less on Model-based Reinforcement Learning. J Cogn Neurosci. 28:985–95.

Seidel EM, Habel U, Kirschner M, Gur RC, Derntl B. (2010). The impact of facial emotional expressions on behavioral tendencies in women and men. J Exp Psychol Hum Percept Perform. 36:500–7.

Swart JC, Fröbose MI, Cook JL, Geurts DE, Frank MJ, Cools R, den Ouden HE. (2017). Catecholaminergic challenge uncovers distinct Pavlovian and instrumental mechanisms of motivated (in)action. Elife. 6. pii: e22169.

Talmi D, Seymour B, Dayan P, Dolan RJ. (2008). Human pavlovian-instrumental transfer. J Neurosci. 28:360–8.

Tottenham N, Tanaka JW, Leon AC, McCarry T, Nurse M, Hare TA, Marcus DJ, Westerlund A, Casey BJ, Nelson C. (2009). The NimStim set of facial expressions: judgments from untrained research participants. Psychiatry Res. 168:242–9.

von Stein A, Sarnthein J. (2000). Different frequencies for different scales of cortical integration: from local gamma to long range alpha/theta synchronization. Int J Psychophysiol. 38:301–13.

Watanabe S. (2013). A Widely Applicable Bayesian Information Criterion. Journal of Machine Learning Research 14:867–897.

Zaki J, Kallman S, Wimmer GE, Ochsner K, Shohamy D. (2016). Social Cognition as Reinforcement Learning: Feedback Modulates Emotion Inference. J Cogn Neurosci. 28:1270–82.

Zaki J, Hennigan K, Weber J, Ochsner KN. (2010). Social cognitive conflict resolution: contributions of domain-general and domain-specific neural systems. J Neurosci. 30:8481–8.

